# Angiotensin-II modulates GABAergic neurotransmission in the mouse substantia nigra

**DOI:** 10.1101/2021.02.02.429274

**Authors:** Maibam R. Singh, Jozsef Vigh, Gregory C. Amberg

## Abstract

GABAergic projections neurons of the substantia nigra reticulata (SNr), through an extensive network of dendritic arbors and axon collaterals, provide robust inhibitory input to neighboring dopaminergic neurons in the substantia nigra compacta (SNc). Angiotensin-II (Ang-II) receptor signaling increases SNc dopaminergic neuronal sensitivity to insult, thus rendering these cells susceptible to dysfunction and destruction. However, the mechanisms by which Ang-II regulates SNc dopaminergic neuronal activity are unclear. Given the complex relationship between SN dopaminergic and GABAergic neurons, we hypothesized that Ang-II could regulate SNc dopaminergic neuronal activity directly and indirectly by modulating SNr GABAergic neurotransmission. Herein, using transgenic mice, slice electrophysiology, and optogenetics, we provide evidence of an AT_1_ receptor-mediated signaling mechanism in SNr GABAergic neurons where Ang-II suppresses electrically-evoked neuronal output by facilitating postsynaptic GABA_A_ receptors and prolonging the action potential duration. Unexpectedly, Ang-II had no discernable effects on the electrical properties of SNc dopaminergic neurons. Also, and indicating a nonlinear relationship between electrical activity and neuronal output, following phasic photoactivation of SNr GABAergic neurons, Ang-II paradoxically enhanced the feedforward inhibitory input to SNc dopaminergic neurons. In sum, our observations describe an increasingly complex and heterogeneous response of the SN to Ang-II by revealing cell-specific responses and nonlinear effects on intranigral GABAergic neurotransmission. Our data further implicate the renin-angiotensin-system as a functionally relevant neuromodulator in the basal ganglia, thus underscoring a need for additional inquiry.

**SIGNIFICANCE STATEMENT:** Angiotensin II (Ang-II) promotes dopamine release in the striatum and, in the substantia nigra compacta (SNc), exacerbates dopaminergic cell loss in animal models of Parkinson’s disease. Despite a potential association with Parkinson’s disease, the effects of Ang-II on neuronal activity in the basal ganglia is unknown. Here we describe a novel AT_1_ receptor-dependent signaling mechanism in GABAergic projection neurons of the SN reticulata (SNr), a major inhibitory regulator of SNc dopaminergic neurons. Specifically, Ang-II suppresses SNr GABAergic neuronal activity, subsequently altering GABAergic modulation of SNc dopaminergic neurons in a nonlinear fashion. Altogether, our data provide the first indication of Ang-II-dependent modulation of GABAergic neurotransmission in the SN, which could impact output from the basal ganglia in health and disease.

## INTRODUCTION

Complementing the systemic cardiovascular renin-angiotensin-system (RAS), the central nervous system contains a fully-formed and independent RAS (Arnold et al., 2013; Labandeira-Garcia et al., 2014; Rocha et al., 2016; Valenzuela et al., 2016; Labandeira-Garcia et al., 2017; Perez-Lloret et al., 2017; Jackson et al., 2018). Increasing evidence suggests that the peptide hormone angiotensin II (Ang-II), a primary RAS effector, contributes to neurodegenerative disorders such as Parkinson’s disease (PD) and Alzheimer’s disease (Arnold et al., 2013; Labandeira-Garcia et al., 2014; Jackson et al., 2018). In animal models of PD, Ang-II-dependent activation of AT_1_ receptors promotes dopaminergic neuronal cell loss in the substantia nigra compacta (SNc) (Grammatopoulos et al., 2007; Sonsalla et al., 2013).

Evidence also suggests that Ang-II evokes dopamine release in the rat striatum, and several studies report altered Ang-II receptor expression levels in tissue samples from Parkinson’s patients (Simonnet and Giorguieff-Chesselet, 1979; Allen et al., 1991; Brown et al., 1996; Ge and Barnes, 1996). Consistent with these observations, dopaminergic and GABAergic neurons in the SN of rodents and primates, including humans, express Ang-II receptors (Garrido-Gil et al., 2013). However, the occurrence of Ang-II-dependent modulation of GABAergic input to SNc dopaminergic neurons remains to be determined.

GABAergic projection neurons in the substantia nigra reticulata (SNr), located ventrolateral to the SNc, through their axon collaterals, provides a major inhibitory input to SNc dopaminergic neurons (Parent, 1990; Mailly et al., 2003; Hikosaka, 2007; Tepper and Lee, 2007). Indeed, more than 70% of the synapses onto SNc dopaminergic neurons are GABAergic (Bolam and Smith, 1990; Tepper and Lee, 2007). This intranigral inhibitory circuit regulates phasic dopaminergic output during behaviors such as reward extinction and conditioned negative association (Pan et al., 2013; Brown et al., 2014). Accordingly, inhibition of SNr GABAergic neurons causes disinhibition and burst firing of SNc dopamine neurons, leading to increased dopamine levels in the striatum and the basal ganglia (Tepper et al., 1995; Tepper and Lee, 2007; Lobb et al., 2011).

Regulation of SNc dopaminergic neurotransmission by SNr GABAergic neurons is well established and contributes to basal ganglia circuit dysfunction (Deniau et al., 2007; Rice et al., 2011). Prior investigations show that Ang-II, via activation of AT_1_ receptors, modulates GABAergic neurotransmission in the anterior hypothalamus, amygdala, and the median preoptic nucleus (Henry et al., 2009; Xing et al., 2009; Hu et al., 2018). Given the strong expression of Ang-II receptors throughout the SN and the predominantly GABAergic control of dopaminergic neurons, Ang-II signaling in SNr GABAergic neurons could potentially modulate intranigral GABAergic and dopaminergic neurotransmission.

Herein, we tested the hypothesis that Ang-II modulates SNr GABAergic neuronal activity, which, in turn, alters GABAergic regulation of SNc dopaminergic neurons. Using a combination of *ex vivo* brain slice electrophysiology and optogenetics, we find that Ang-II, via AT1 receptors, acutely suppresses electrically evoked action potential (AP) firing of SNr GABAergic neurons. This effect depended on enhanced postsynaptic GABA_A_-receptor activity and prolonged action potential duration (APD). Paradoxically, upon phasic photoactivation of SNr GABAergic neurons, Ang-II enhanced the feedforward inhibition of SNc dopaminergic neurons. This unexpected finding suggests that Ang-II’s effect on postsynaptic SN dopaminergic neurons is nonlinear during synchronous GABAergic neuronal activity.

In sum, our data provide strong evidence of Ang-II signaling in SN GABAergic neurons and illustrate the complex heterogeneity of the ensuing neuronal responses. Further, our observations reveal the intranigral microcircuitry as a potential target for modifying the basal ganglia output.

## MATERIALS AND METHODS

### Animals

We bred the following two mouse strains from The Jackson Laboratory to generate a tdTomato reporter mice for dopaminergic neurons (Fig. 1*A*): Cre-dependent tdTomato reporter mice [B6.Cg-*Gt(Cooper et al.)26Sor^tm9(CAG-tdTomato)Hze^*/J, stock #007909], and tyrosine hydroxylase (TH) promotor-driven Cre expression [B6.Cg-*7630403G23Rik^Tg(Th-cre)1Tmd^*/J, stock #008601]. In order to focally stimulate SNr GABAergic neurons in *ex-vivo* brain slices, we used transgenic mice expressing ChR2 fused to YFP under the control of mouse thymus cell antigen 1 (*Thy 1*) promoter (stock #007612, Jackson laboratory), which specifically express ChR2 in SNr GABA neurons, but not in SN dopaminergic neurons. All mice used for the study were between 4-8 weeks old. To detect transgene and floxed alleles, we utilized a genotyping service (Transnetyx).

**Figure 1.**
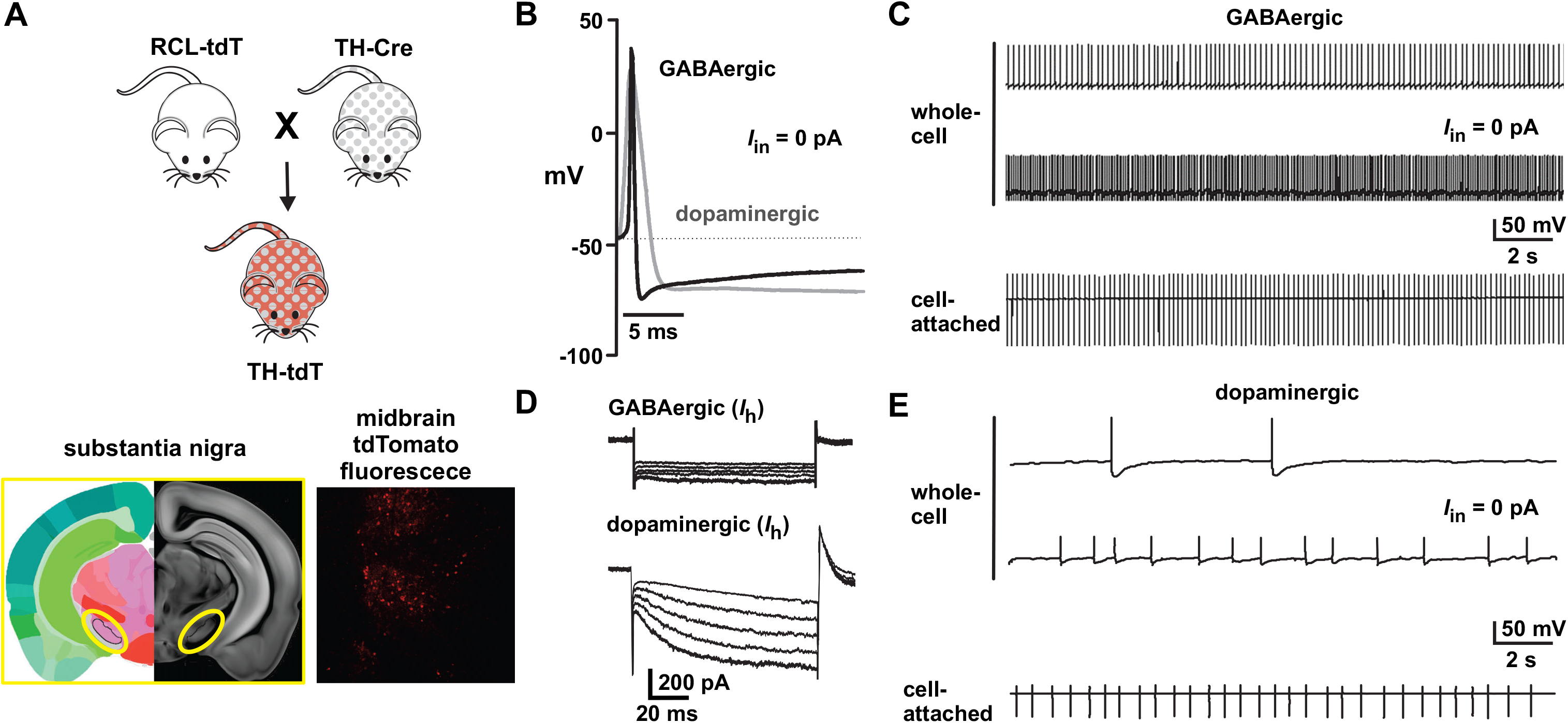
Electrophysiological characteristics of GABAergic and dopaminergic neurons in the mouse substantia nigra. ***A***, RCL-tdT Cre reporter mice crossed with tyrosine hydroxylase (TH) promotor-dependent Cre expressing produce mice with td-Tomato expression (*red fluorescence*) restricted to TH expressing dopaminergic neurons. SNr GABAergic neurons were differentiated from SNc dopaminergic by lack of td-Tomato fluorescence and distinct electrophysiological profile. ***B***, SNr GABAergic neurons have a narrower (< 1.5 ms) action potential width than SNc dopaminergic neurons (> 3 ms). ***C***, Representative sustained high-frequency firing of SNr GABAergic neurons recorded in whole-cell (*top*) and cell-attached configuration (*bottom*). ***D,*** SNr GABAergic neurons, in contrast to SNc dopaminergic neurons, show little to no *I*_h_ in response to a series of hyperpolarizing pre pulses (−70 mV to - 140 mV). ***E***, Representative low-frequency, slow, and irregular pacemaker-like firing in SNc dopaminergic neurons.

### Animal care and euthanasia

Mice received *ad libitum* access to standard chow and tap water while housed individually or in groups of less than four in a temperature and humidity-controlled room set on a 12-h light/dark cycle. On the day of experimentation, adult male and female mice were deeply anesthetized under isoflurane, decapitated, and the brains were removed and placed into an ice-cold artificial cerebral spinal fluid (aCSF) containing (mM): 126 NaCl, 2.5 KCl, 1.2 MgCl_2_, 1.4 NaH_2_PO_4_, 25 NaHCO_3_, 11 D-Glucose, 2.4 CaCl_2_, pH = 7.4, osmolarity = 310 mOsm) bubbled with 95% O_2_/5% CO_2_. All procedures, including euthanasia, were performed according to institutional guidelines and approved by the Institutional Animal Care and Use Committee of Colorado State University.

### Slice electrophysiology

We used midbrain coronal slices to minimize the influence of presynaptic connections and carry out electrophysiological recordings in relative isolation. Coronal slices (≈ 240 μM thick) containing the substantia nigra reticulata (SNr) and compacta (SNc) were cut (Leica VT1200S; Leica Microsystems), transferred to a holding chamber, and incubated for 30-60 min at 35°C in aCSF supplemented with MK-801 (100 μM), and stored at 21°C until used for experimentation. To record from cells, we transferred individual slices to a chamber continuously perfused with 21-22°C aCSF at a rate of 2-3 ml/min. We identified and differentiated nigral GABAergic and dopaminergic based on their well-established electrophysiological features (described below) as well as expressed fluorescence in the respective transgenic lines: TdTomato for dopaminergic neurons (TH-Cre-TdTomato) and YFP for GABAergic neurons (Thy1-ChR2-YFP).

Borosilicate glass pipettes, with resistances of 3-5 MΩ, were fabricated on a laser micropipette puller (Model P-2000; Sutter Instrument). Whole-cell current-clamp recordings of evoked spike activity used a potassium gluconate-based intracellular solution composed of (mM): 120 K-gluconate, 20 KCl, 10 HEPES, 0.2 EGTA, 2 MgCl_2_, 10 phosphocreatine, 2 Mg-ATP, 0.3 Na-GTP, pH = 7.5 adjusted with KOH, osmolarity = 290-295 mOsm). Pipettes for recording spontaneous postsynaptic currents (PSCs) and miniature inhibitory PSCs (mIPSCs) contained a high chloride potassium methylsulfate-based solution composed of (mM): 57.5 K-methyl sulfate, 57.5 KCl, 20 NaCl, 1.5 MgCl_2_, 5 HEPES, 0.1 EGTA, 10 phosphocreatine, 2 Mg-ATP, 0.3 Na-GTP, pH = 7.5 adjusted with KOH, osmolarity = 290-295 mOsm).

We visualized cells with a 40X water immersion objective on an upright microscope (Zeiss) equipped with Dodt gradient contrast and collected data with an EPC-10 USB patch-clamp amplifier controlled with PatchMaster software (v.2.30; HEKA Electronik). Data were low-pass filtered at 10 kHz and sampled at 50 kHz. We compensated for fast and slow capacitive transients, series resistance, and only used recordings with stable series resistances < 20 MΩ. To ensure the fidelity of our measurements, we continuously monitored electrophysiological parameters, including series resistance, leak, and membrane voltage. Cells with unstable recording parameters, such as leak change of > 20% and high series resistance, were flagged and not used. GABAergic SNr neurons were electrophysiologically identified and distinguished from dopaminergic neurons by their well-characterized electrophysiological profile (Yung et al., 1991; Richards et al., 1997; Pan et al., 2013).

After gaining whole-cell access, cells were held at −65 mV (corrected for a liquid junction potential of 5 mV (Barry, 1994)). Following stabilization, cells were hyperpolarized stepwise from −65 mV to −140 mV to measure I_h_ and provide an initial characterization of the cell as either GABAergic or dopaminergic. Next, evoked APs were recorded in response to 2 s long current injections, increasing stepwise from −50 pA to 150 pA, for at least 5 mins. After obtaining a series of stable recordings with aCSF (control), we superfused cells with aCSF supplemented with Ang-II (500 nM) for a minimum of 5 mins and recorded evoked APs for ≥ 15 mins. To conclude the experiment, Ang-II was washed out by superfusion with standard aCSF for at least 5 mins, after which a series of final recordings were recorded. For experiments with the AT_1_ receptor antagonist losartan (1 μM) and the GABA_A_R antagonist picrotoxin (1 μM), each drug was superfused alone or with Ang-II. For these recordings, we used the same experimental design as used for Ang-II alone.

We recorded spontaneous whole-cell outward PSCs with a high chloride (57.5 mM) internal solution (described above) at a liquid junction potential-corrected setting of −70 mV. We did not correct for leak and discarded any cell with leak > ± 100 pA or a change in leak > ± 50 pA during the length of the recording. Ang-II was either bath perfused for 3-5 mins or puffed using a 1-3 MΩ glass pipette positioned ahead of the recording pipette and in the direction of laminar flow with Picospritzer III (Parker, Cleveland, OH). To isolate and record spontaneous mIPSCs at a holding potential of - 70 mV, we blocked excitatory synaptic transmission using a cocktail of drugs: the voltage-dependent sodium blocker tetrodotoxin (TTX; 500 nM), the AMPA/kainate receptor antagonist CNQX (1 μM), the NMDA receptor antagonist MK-801 (1 μM), and the nicotinic receptor antagonist hexamethonium bromide (100 μM). IPSC’s were confirmed to be mediated by GABA_A_ receptors with picrotoxin (100 μM).

### Photostimulation and slice electrophysiology

To generate light-evoked EPSPs from ChR2 expressing SNr GABA neurons, we applied three 100 ms long light pulses with an interstimulus interval of 2 s using a 470 nM LED (Thorlabs) driven by a LEDD1B driver (Thorlabs). Intrabusrt EPSPs and spikes from three light-evoked responses were averaged with a minimum of 3-5 recordings from each group. We used a K-gluconate based internal solution (described above) for recording light-evoked EPSPs and IPSCs. IPSCs in SNc dopaminergic neurons were recorded in the presence of excitatory synaptic blockers and identified as outward current deflections in response to photostimulation of SNr GABA neurons. For analysis, we included only SNc dopaminergic neurons with outward current deflections in response to photostimulation of SNr GABAergic neurons. Similarly, neurons with light-evoked excitatory postsynaptic potentials (EPSPs) were classified as GABAergic, whereas neurons showing light-evoked IPSPs were classified as dopaminergic. EPSPs and IPSPs were recorded in current-clamp mode at −65 mV (adjusted for junction potential) using the K-gluconate based internal solution.

### Drugs

We purchased CNQX, MK-801, Ang-II, losartan, and picrotoxin from Sigma; and TTX and PD123319 from Tocris Biosciences. Drugs were prepared immediately before use in either distilled water (Ang-II, TTX, PD123319, and losartan) or DMSO (CNQX, MK-801, and picrotoxin) and diluted in aCSF to achieve the desired concentration. We used 500 nM Ang-II for our experiments, a high working concentration for a ligand-receptor pair with a Kd less than 5 nM. However, we used a high Ang-II concentration to ensure adequate slice penetration of bath-applied Ang-II.

### Experimental Design and Statistical analyses

Data were analyzed using Axograph X and GraphPad Prism (v.8) software. We used one-way, two-way ANOVA’s, mixed-effect analyses, and paired student t-test as indicated. Individual points in the figures represent data from a single cell. We performed only one experiment per brain slice and obtained no more than three recordings per mouse. We used male and female mice for all experiments. However, a sex-difference analysis revealed nothing of significance; thus, we pooled our data. Averaged data are presented as the mean ± SEM unless otherwise specified; *n* = number of cells recorded from, and significance was defined as *p* < 0.05 unless otherwise indicated.

## RESULTS

To test our hypothesis that Ang-II regulates the activity of GABAergic projection neurons in the mouse SNr, we formulated the following requisite experimental criteria: 1) exogenous Ang-II must alter the AP firing characteristics of positively identified SNr GABAergic neurons in *ex vivo* brain slices; 2) Ang-II must promote changes in the electrophysiological properties of SNr GABAergic neurons by mechanisms consistent with the observed changes in AP firing behavior; 3) the observed effects of Ang-II must be sensitive to pharmacological blockade of cognate Ang-II receptors and; 4) Ang-II should suppress light-evoked EPSPs in ChR2 expressing SNr GABAergic neurons and; 5) Ang-II must modulate GABAergic input onto postsynaptic SNc dopaminergic neurons.

### Ang-II suppresses evoked action potentials in SNr GABAergic neurons

We performed *ex-vivo* whole-cell electrophysiology on freshly prepared coronal midbrain slices from TdTomato dopaminergic neuron reporter mouse to investigate the effects of Ang-II on GABAergic projection neurons in the SNr. To begin, we used whole-cell current-clamp to record spontaneous and depolarization-evoked action potentials. As a means to provisionally identify and distinguish GABAergic neurons from dopaminergic neurons, we selected cells by their lack of tdTomato fluorescence and neuroanatomical features (Fig. 1*A*; see Materials and Methods). After obtaining electrophysiological access, we confirmed or contested cell identities using the well-characterized electrophysiological profiles of GABAergic and dopaminergic neurons (Franklin and Paxinos, 2001; Lacey et al., 1989; Tepper et al., 1995; Richards et al., 1997; Paxinos and Franklin, 2004). Thus, criteria used to categorize cells as GABAergic included: 1, an apparent absence of tdTomato fluorescence; 2, the presence of sustained high-frequency spontaneous AP firing (> 10 Hz); 3, an action potential width of < 2 ms; 4, little or no hyperpolarization currents; and 5, minimal adaptation to injected depolarizing currents (Fig. *1B - D*). Alternatively, we identified neurons as dopaminergic based on their cell size, anatomical location, detectable tdTomato fluorescence, slow pacemaker-like AP firing, action potential durations > 2 ms, robust hyperpolarization currents, and pronounced adaptation to injected depolarizing currents.

To achieve a relatively controlled measure of AP activity, we used an evoked AP protocol consisting of 2 s current injections increasing incrementally from −50 pA to +150 pA. In cells identified as GABAergic, following control recordings in unsupplemented aCSF, bath-applied Ang-II (0.5 μM) decreased evoked AP firing during current injections of 50, 100, and 150 pA (Fig. *2A & B*, *n* = 17, *p* = 0.003 for 50 pA, *p* < 0.001 for 100 pA and 150 pA, F_(2,32)_= 9.521; mixed-effect analysis). In 10 out of 17 cells, the effect of Ang-II on evoked AP firing was partially reversible with washout. Illustrating the degree of AP firing suppression, Ang-II increased the mean interspike interval by approximately two-fold (Fig. 2*C*; n = 17, F_(2,29)_= 5.976, *p* = 0.005; mixed-effect analysis). Additionally, Ang-II also increased the variability (i.e., irregularity) of the interspike intervals (Fig. 2*D*; *n* = 16, F_(2,28)_ = 5.488, *p* = 0.01; mixed-effect analysis).

**Figure 2.**
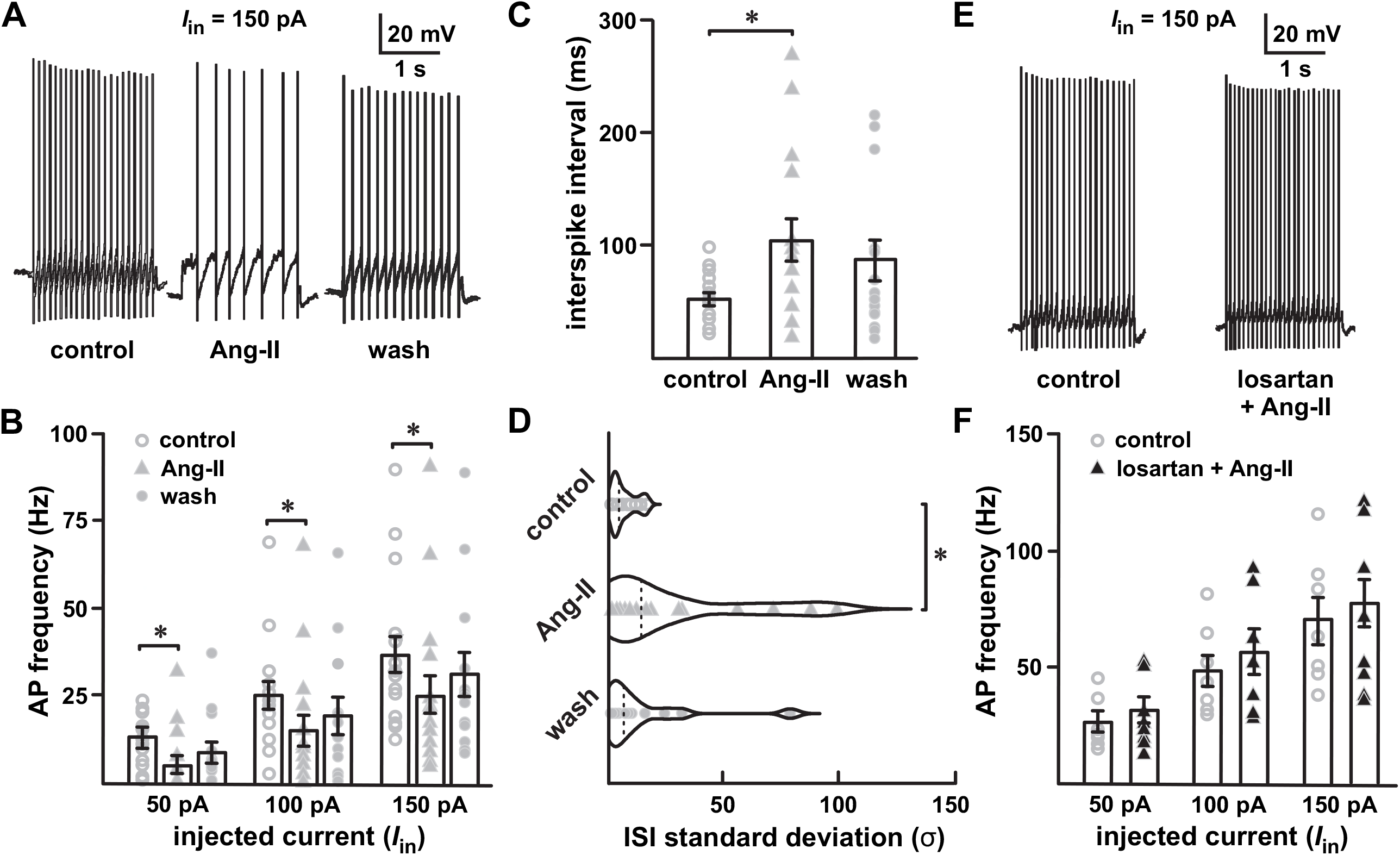
Angiotensin-II decreases SNr GABAergic projection neuron spike firing. ***A***, Evoked action potentials in response to 150 pA current injection. ***B,*** Individual scatter plot of evoked spike frequency in the SNr GABAergic projection neurons in response to 50 pA, 100 pA, and 150 pA current injections. Ang-II significantly decreased evoked firing in all stimulus levels, 50 pA-150 pA (*** *p* < 0.001, two-way repeated-measures ANOVA). In 10 out of 17 cells, washout partially reversed the effect of Ang-II. ***C***, Ang-II increased the SNr GABAergic interspike interval (ISI; \*\**p* = 0.005, mixed effect analysis) and (***D***) increased the irregularity of firing as quantified by the ISI standard deviation (* *p* = 0.01, mixed effect analysis)***. E & F,*** The AT_1_-R specific blocker losartan (1 μM) abolished the suppression of evoked SNr GABAergic spike firing by Ang-II.

Indicative of AT_1_ receptor involvement, preincubation with losartan (1 μM) abolished decreased evoked AP firing following Ang-II application (Fig. 2*E* & *F*). To test for potential contributions by Ang-II type 2 receptors (AT_2_Rs), we used the specific AT_2_R antagonist PD123319. In contrast to AT_1_ receptor blockade with losartan, PD123319 (1 μM) did not block Ang-II-dependent suppression of SNr GABAergic neuronal activity (*n* = 5, F_(1,4)_ = 5.896, *p* = 0.072, Repeated measures Two-way ANOVA; data not shown). From these data, we conclude that Ang-II decreases evoked AP firing in SNr GABAergic neurons via AT_1_ receptor signaling. Note that we collected comparable data on identified SNc dopaminergic neurons (data not shown). However, the well-described adaptive responses of dopaminergic neurons to depolarization, and eventual depolarization block, precluded meaningful interpretation of these data (Lacey et al., 1989; Richards et al., 1997).

### Ang-II prolongs action potential durations in SNr GABAergic neurons

Sustained spontaneous high-frequency AP firing (10 – 15 Hz, *in vitro*) is a striking characteristic of SNr GABAergic neurons (Lacey et al., 1989; Richards et al., 1997; Atherton and Bevan, 2005; Zhou and Lee, 2011). The high-frequency firing in these cells is autonomously generated and involves multiple ion channels from more than five families (Atherton and Bevan, 2005; Zhou et al., 2008; Seutin and Engel, 2010; Ding et al., 2011). We found that in SNr GABAergic neurons, Ang-II prolonged the duration of high-frequency APs in a largely reversible fashion (Figs. 3 & 4). Indeed, Ang-II increased the half-width of both evoked (Fig. 3*C*; *n* = 17; *F*_(2,29)_ = 5.440; *p* = 0.010; mixed-effect analysis) and spontaneous APs (Fig. 4*C*; *n* = 5; *F*_(2,8)_ = 9.481; *p* = 0.008; one-way repeated measures ANOVA). The effects of Ang-II on AP duration were prevented by preincubation with losartan (1 μM; Fig. 4*B*).

**Figure 3.**
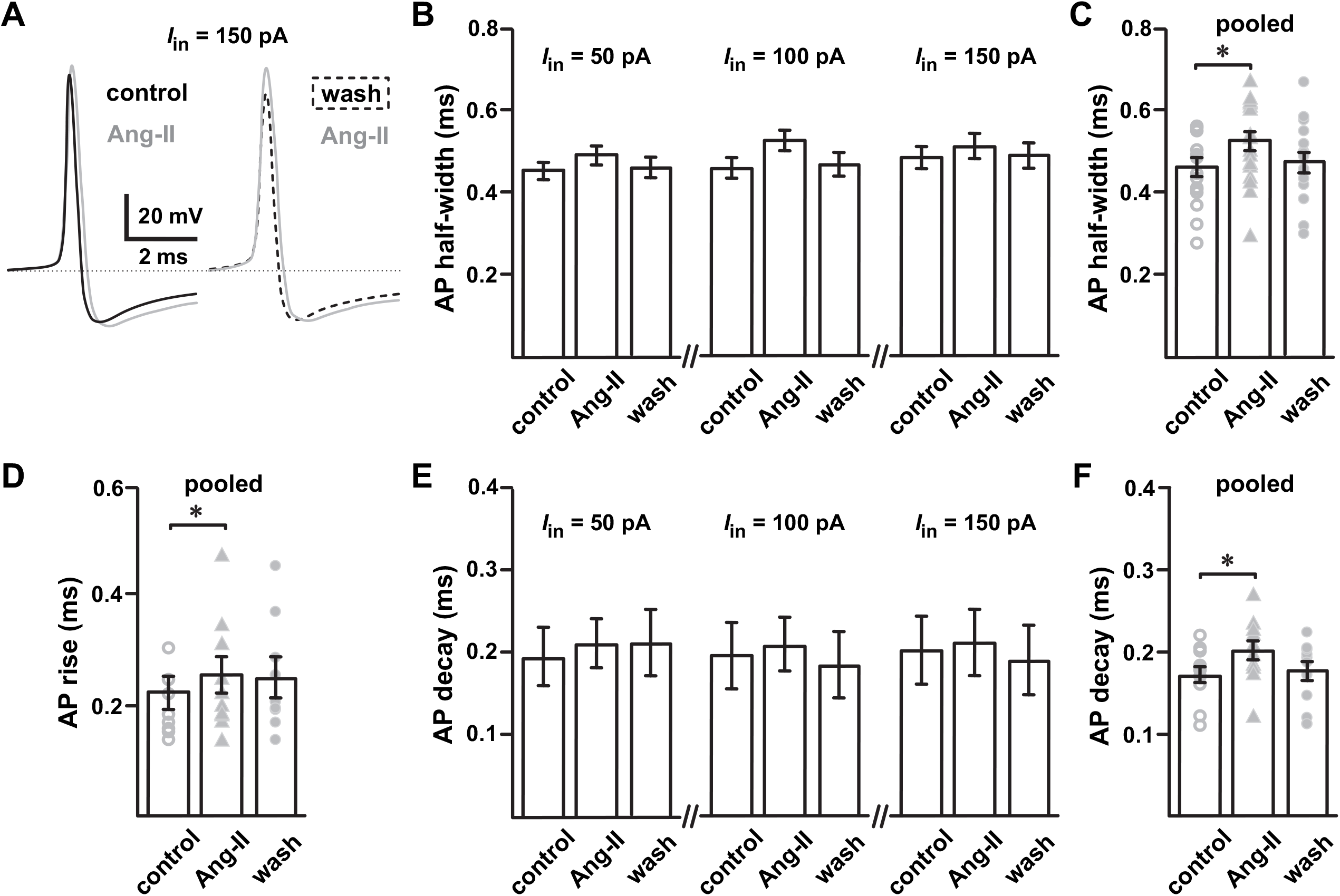
Ang-II slows the action potential kinetics of SNr GABAergic neurons. ***A***, Representative AP waveforms of SNr projection neurons showing that Ang-II reversibly slows AP kinetics. ***B & C,*** Ang-II increased the AP half-width of SNr GABAergic neurons (* *p* = 0.010, mixed-effect analysis). ***D***, Ang-II slowed the rise of APs in SNr GABAergic neurons (** *p* = 0.005, repeated measures one-way ANOVA). ***E & F,*** Ang-II slowed the decay of APs in SNr GABAergic neurons (^**^*p* = 0.004, repeated measure one-way ANOVA).

**Figure 4.**
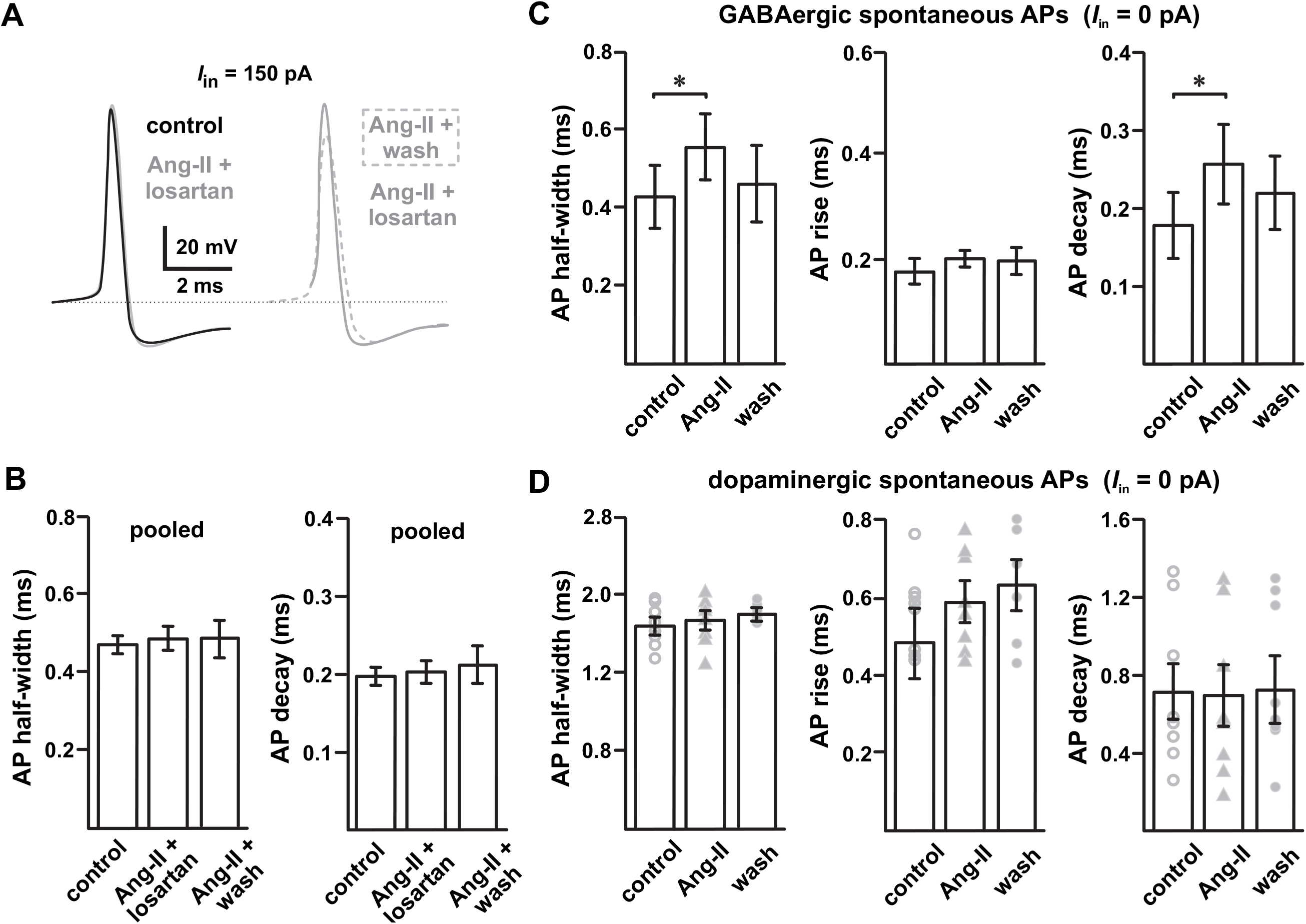
Disparate effect of Ang-II on SN GABAergic and dopaminergic neurons. ***A,*** Losartan blocks Ang-II mediated increase in AP duration in SNr GABAergic neurons; losartan washout with continued Ang-II perfusion prolonged the AP duration. ***B***, Summary data showing that losartan (1 μM) blocks Ang-II mediated increase in AP half-width and decay. ***C,*** Ang-II increased the duration of SNr GABAergic neuron spontaneous APs (* *p* = 0.04, AP half-width; * *p* = 0.015, AP decay; repeated measures one-way ANOVA). ***D,*** Unlike SNr GABAergic neurons, Ang-II had no noticeable effect on the AP kinetics of SNc dopaminergic neurons.

Further analysis of evoked APs in Ang-II-responsive SNr GABAergic neurons (i.e., those cells showing AP prolongation with Ang-II; 11 of 17 cells) revealed slower AP kinetics after Ang-II application. Specifically, Ang-II slowed evoked AP rise-time (Fig. 3*D*; *n* = 9; *F*_(2,16)_ = 7.683; *p* = 0.005; repeated measures one-way ANOVA) and slowed AP decay (Fig. 3*F*; *n* = 10; *F*_(2,18)_ = 7.102; *p* = 0.005; repeated measure one-way ANOVA). Although no apparent effect on spontaneous AP rise times was evident, Ang-II also slowed the decay of spontaneous APs in GABAergic neurons (Fig. 4*C*; *n* = 5; *F*_(2,8)_ = 9.207; *p* = 0.015; repeated measures one-way ANOVA). Interestingly, and contrasting SNr GABAergic neurons, cells identified as dopaminergic had no observable changes in AP duration or kinetics in response to Ang-II (Fig. 4*D*). These data suggest that AT_1_ receptor activation by Ang-II prolongs AP durations of SNr GABAergic but not SNc dopaminergic neurons. Further, the observed suppression of AP firing of SNr GABAergic neurons following Ang-II administration could arise, at least in part, as a consequence of AP prolongation due to a slowed rise and decay kinetics.

### Ang-II potentiates postsynaptic GABA_A_ receptors in SNr GABAergic neurons

Striatal GABAergic medium-sized spiny neurons provide tonic inhibitory input to the SNr GABAergic neurons (Dray, 1979; Graybiel and Ragsdale, 1979; McGeer et al., 1984; Smith and Bolam, 1990). These GABAergic neurons in the SNr express mostly GABA_A_ receptors but few GABA_B_ receptors. Indeed, electrical stimulation of the striatum produces short-term inhibition of SNr GABAergic neurons that is blocked by the GABA_A_ receptor antagonists such as picrotoxin (Precht and Yoshida, 1971; Dray, 1979; Wallmichrath and Szabo, 2002). To test if the observed suppression of SNr GABAergic neuronal activity by Ang-II involves GABA_A_ receptors (see Fig. 2), we replicated our evoked AP experiments in the presence of GABA_A_ receptor antagonist picrotoxin. Suggesting GABA_A_ receptor involvement, preincubation with picrotoxin (1 μM) attenuated Ang-II-dependent suppression of GABAergic neuronal firing (Fig. 5*B*). Interestingly, picrotoxin did not completely prevent Ang-II from prolonging the AP duration and slowing the AP rise time (Fig. 5*C*; *n* = 5, *p* = 0.103; Student’s paired t-test), decay (Fig. 5*C*; *n* = 5, *p* = 0.101; student’s paired t-test) and half-width (Fig. 5*C*; *n =* 5, *p =* 0.119) in these cells. These data suggest that Ang-II potentially suppresses the excitability of SNr GABAergic neurons by two independent mechanisms (slowed AP kinetics and potentiation of GABA_A_ receptors).

**Figure 5.**
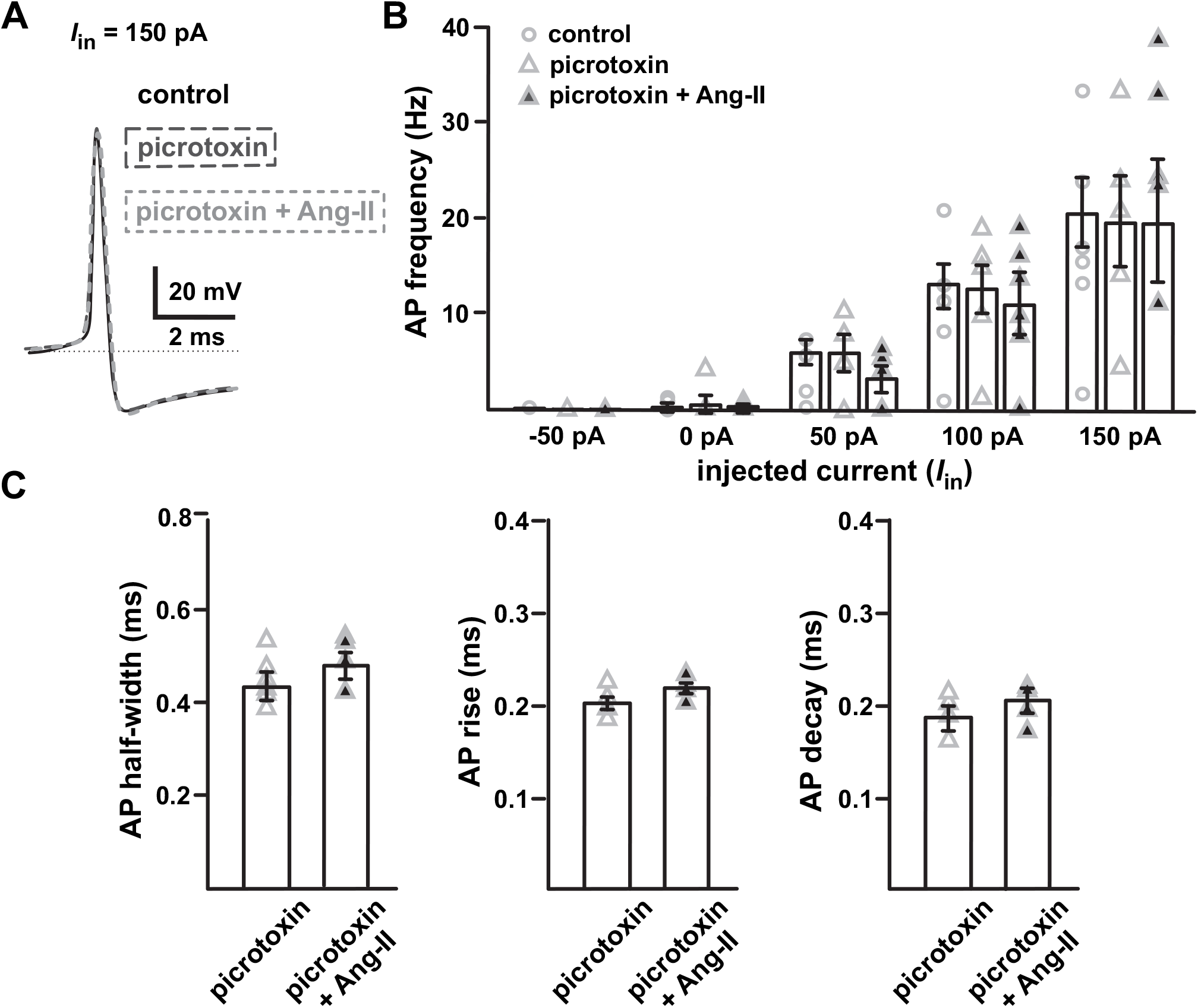
GABA_A_ receptor blockade prevents Ang-II mediated suppression of evoked spike firing in the SNr GABAergic projection neurons. *A*, Overlaid spike waveforms for control conditions, in the presence of the GABA_A_ receptor antagonist picrotoxin (1 μM), and in the presence of picrotoxin plus Ang-II. *B,* Summary plot showing evoked spike firing of SNr GABAergic neurons under control conditions, in the presence of picrotoxin (1 μM), and in the presence of picrotoxin plus Ang-II. *C,* GABA_A_ receptor blockade with picrotoxin attenuates Ang-II mediated increases in SNr GABAergic neuron AP durations.

To further examine the effects of Ang-II-dependent modulation of GABA_A_ receptor activity in SNr GABAergic projection neurons, we recorded spontaneous postsynaptic currents (PSCs) (Fig. 6). Although Ang II did not significantly alter the frequency or average amplitude of spontaneous PSCs in these cells (Fig. 6*B*), we did observe a roughly two-fold increased incidence of high-amplitude spontaneous PSCs, which are 3X the mean amplitude (≥ −150 pA) (Fig. 6*B*; *n* = 9; *p* = 0.012; paired Student’s *t*-test). Finally, we replicated our spontaneous PSC experiments in the presence of AP and excitatory synaptic blocker cocktail (see Materials and Methods) to confirm that the Ang-II-dependent increases in PSCs are mediated postsynaptically. Synaptic blockade drastically reduced the overall activity in our recordings to reveal the presence of spontaneous miniature IPSCs (mIPSCs; Fig. 7*A*). Similar to our experiments in the absence of synaptic blockers (Fig. 6), Ang-II significantly increased the observed incidence of high amplitude mIPSCs (≥ 3X the mean amplitude; ≥ −121.86 pA), by approximately 2.5-fold without a significant change in the average mIPSC amplitude or frequency (Fig. 7*A & B*; *n* = 12; *p* = 0.001; paired Student’s t-test). Notably, picrotoxin (1 μM) abolished the Ang-II-dependent increase in high-amplitude mIPSCs in all cells (Fig. 7). We repeated the experiment using a Picospritzer III (Parker) to provide a localized puff of Ang-II near the patched cell; however, we did not observe a noticeable change in high amplitude mIPSC’s (data not shown). In contrast, when Ang-II was reintroduced into the bath, the increase in high amplitude IPSCs occurred as before (data not shown). This observation suggests that the GABA_A_ receptors affected by Ang-II are likely located in distant dendritic terminals away from the soma and are embedded deep within the slice and cannot be reached with a topical puff of Ang-II. From these data, we conclude that Ang-II potentiates postsynaptic GABA_A_ receptors in SNr GABAergic neurons, supporting our initial observation of suppressed excitability of nigral neurons by Ang-II (see Fig. 2).

**Figure 6.**
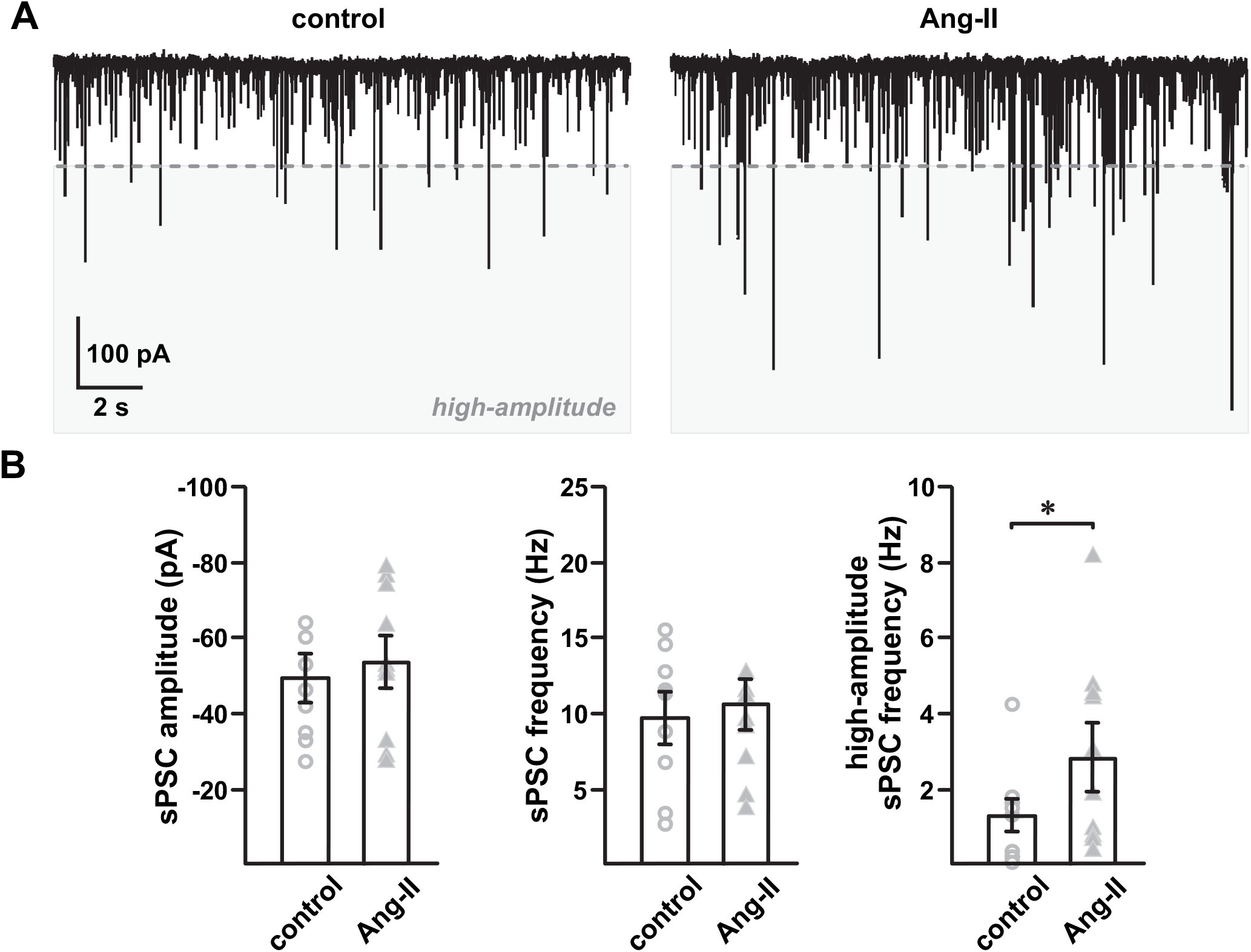
Ang-II increases SNr GABAergic neuron spontaneous postsynaptic currents. ***A***, At a holding potential of - 65 mV (junction potential corrected), spontaneous postsynaptic currents (PSCs) were recorded with a high chloride (57.5 nM) internal pipette solution before and after 5 min of Ang-II (500 nM) exposure. ***B***, Ang-II did not markedly change the average PSC amplitude (*p* = 0.211, paired Student’s t-test) and frequency (*p* = 0.466, paired Student’s t-test) of spontaneous postsynaptic Cl^−^ currents. However, Ang-II significantly increased the observed incidence of high-amplitude spontaneous postsynaptic currents (*right*; * *p* = 0.012; paired Student’s t-test). High-amplitude spontaneous PSCs, demarcated by the dashed line, were defined a priori as currents with an amplitude ≥ three times the mean current amplitude under control conditions.

**Figure 7.**
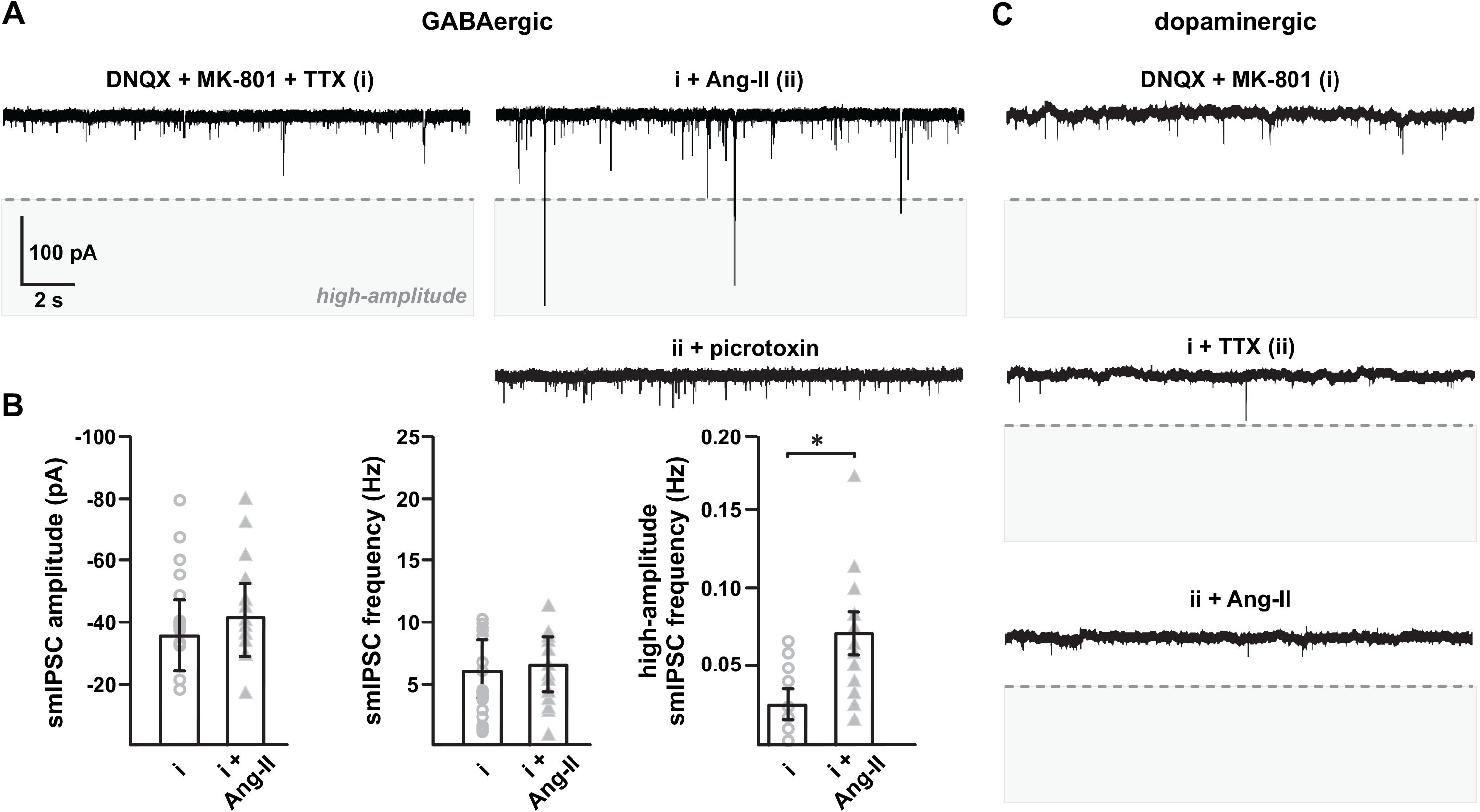
Ang-II increases picrotoxin-sensitive spontaneous miniature IPSCs in SNr GABAergic neurons. Spontaneous mIPSCs recorded from SNr GABAergic and SNc dopaminergic neurons in the presence of excitatory synaptic blockers (DNQX, MK-801; 1 μM)) and TTX (500 nM). ***A***, Representative traces showing Ang-II-mediated enhancement of high-amplitude postsynaptic GABA_A_ receptor currents. High-amplitude spontaneous PSCs, demarcated by the dashed line, were defined a priori as currents with an amplitude ≥ three times the mean current amplitude recorded in the presence of excitatory synaptic blockers and TTX. ***B***, Summary data showing Ang-II-mediated increases in high-amplitude mIPSCs (*p* = 0.019, one-way repeated measures ANOVA), but not in average mIPSC amplitude and frequency. ***C,*** In contrast to SNr GABAergic neurons, Ang-II had no detectable effect on spontaneous mIPSCs in SNc dopaminergic neurons.

### GABA_A_ receptors in SNc dopaminergic neurons are not affected by Ang-II

We also recorded spontaneous mIPSC’s from SNc dopaminergic neurons and found minimal background mIPSC’s compared to SNr GABA neurons (Fig 7*B & C*). Further, Ang-II did not produce a detectable change in the spontaneous mIPSC’s of SNc dopamine neurons (Fig 7*C*). This absence of effect suggests that in the basal state, in contrast to SNr GABA neurons, Ang-II has very little or no direct postsynaptic effect on the GABAergic activity in SNc dopaminergic neurons.

### Ang-II increases GABAergic input to SNc dopaminergic neurons

In our previous experiments, Ang-II suppressed electrically evoked spike output in individual GABAergic neurons. However, the activity of a single neuron is likely not to be representative of the population. Further, the combined input from numerous neurons encode and sum to form the output characteristic of a neuronal population. Accordingly, to induce phasic synchronous neural activity in SNr GABAergic neurons, we used Thy1-ChR2-YFP mice, which expresses ChR2 in SNr GABAergic neurons but not in SNc dopaminergic neurons. To confirm if Ang-II suppression of electrically-evoked firing of individual SNr GABAergic neurons occurs during phasic photoactivation of the population, we recorded EPSPs (Fig 8*A*) from SNr GABAergic neurons during photoactivation with a 100 ms long 470 nm light pulse. Consistent with our earlier observation, Ang-II (0.5 μM) decreased the number of intraburst spikes in 4 out of 6 SNr GABAergic neurons (Fig 8*B & C*). This observation is consistent with our other data showing negative modulatory effects of Ang-II on electrically evoked spike output of SNr GABAergic neurons.

**Figure 8.**
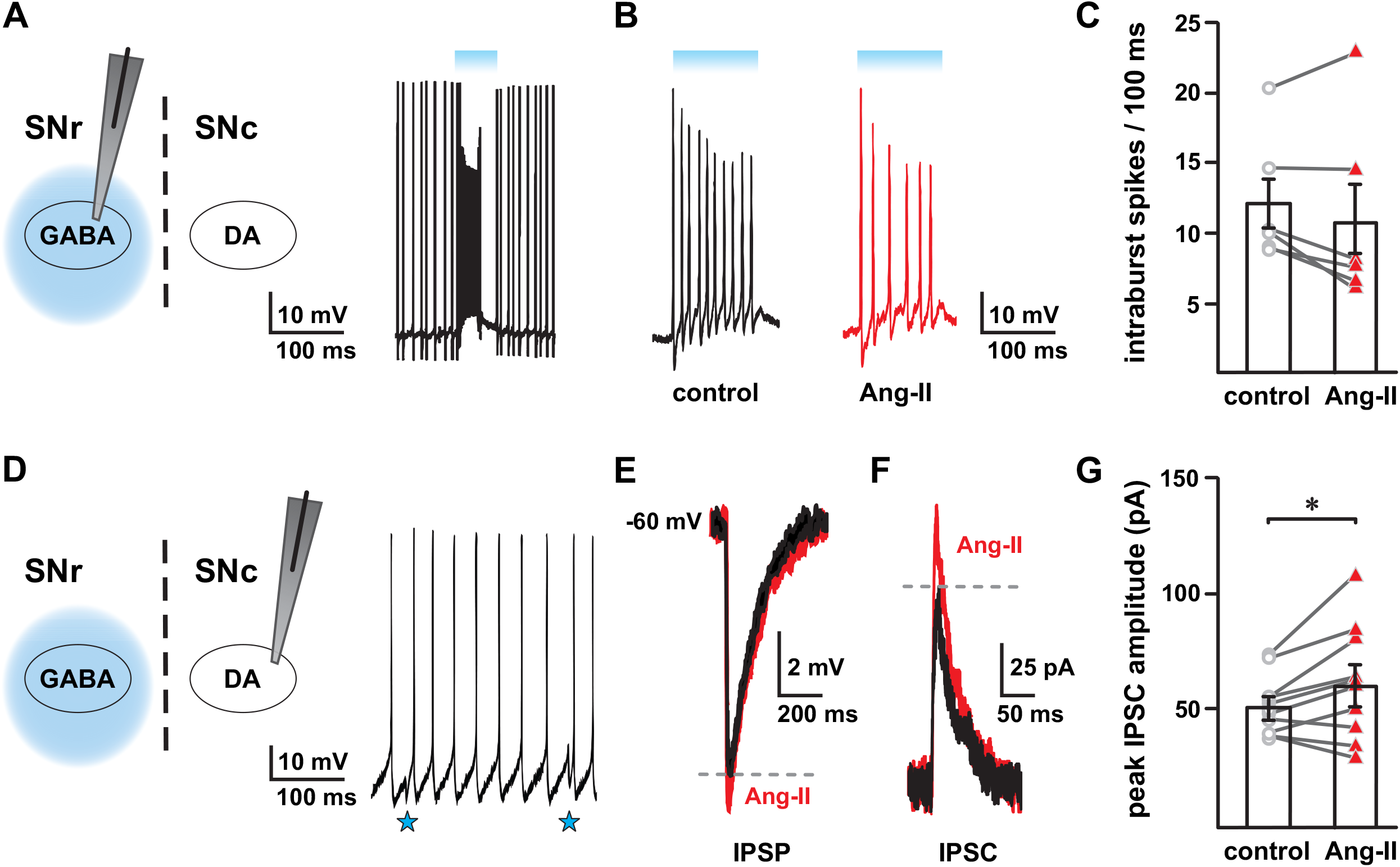
Ang-II modulates SN GABAergic neurotransmission. ***A,*** Light-evoked EPSPs (bursts) in SNr GABAergic neurons in response to a 100 ms long light pulse. ***B,*** Ang-II decreased light-evoked intraburst spikes in SNr GABAergic neurons; intraburst spikes shown are enlarged from *panel A*. ***C,*** Summary data showing the effect of Ang-II on intraburst spikes (EPSPs) in SNr GABAergic neurons in response to 100 ms long light pulse. In 4 out of 6 SNr GABAergic neurons, Ang-II decreased intraburst spikes by ~ 25%. ***D,*** IPSPs recorded from SNc dopaminergic neurons in response to photostimulation of SNr GABAergic neurons, as shown by downward deflection of membrane potential (*blue stars*). Ang-II enhanced light-evoked inhibitory input onto SNc dopaminergic neurons as shown by an increase in the amplitude of light-evoked IPSPs (***E***) and IPSCs (***F & G***; * *p* = 0.049, paired Student’s t-test).

SNr GABAergic neurons provide robust, monosynaptic inhibition of SNc dopaminergic neurons (Tepper et al., 1995; Tepper and Lee, 2007; Brazhnik et al., 2008; Pan et al., 2013). As an alternative approach to measure the effect of Ang-II on the output of SNr GABAergic neurons, we monitored the activity of postsynaptic dopaminergic neurons of the SNc by recording IPSCs and IPSPs from these cells during phasic (100 ms) photoactivation of SNr GABAergic neurons. In our other experiments, Ang-II suppressed electrical and light-evoked activity of SNr GABAergic neurons. Therefore, we expected that Ang-II would disinhibit postsynaptic SNc dopaminergic neurons due to decreased GABAergic input. Unexpectedly, we found that Ang-II produced an ~ 18% increase in the amplitude of light-evoked IPSC’s in SNc dopaminergic neurons (Fig 8*F* & *G*; *n* = 10; *p* = 0.049; paired Student’s t-test). Further, Ang-II also produced a proportional increase in the amplitude of light-evoked IPSP amplitude of these cells (Fig 8*E*).

Together, these data indicate that Ang-II-mediated suppression of SNr GABAergic neurons, as shown in our preceding experiments, does not have a corresponding disinhibitory effect on postsynaptic SNc dopaminergic neurons during synchronous population activity. Rather, Ang-II paradoxically strengthens feedforward inhibition of SNc dopaminergic neurons, suggesting a nonlinear effect of Ang-II on cellular activity and population output of SNr GABAergic neurons.

## DISCUSSION

In this study, we tested the hypothesis that Ang-II signaling in SN GABAergic neurons exerts an acute modulatory influence on mouse SNr GABAergic projection neurons and regulates inhibitory feedforward input to SNc dopaminergic neurons.

### Canonical AT1 receptor signaling acutely and reversibly suppresses evoked action potentials in SNr GABAergic neurons

In neurons identified as GABAergic, we found that acute application of Ang-II decreased evoked AP firing frequencies. We found that the effects of Ang-II on SNr GABAergic neurons were sensitive to AT_1_ receptor blockade by the specific antagonist losartan. This finding is consistent with AT_1_R-mediated modulation of GABAergic neurotransmission in the amygdala and hypothalamus. Consistent with a dynamic regulatory mechanism, Ang-II suppression of SNr GABAergic AP firing was largely reversible upon washout.

In contrast to the blockade of AT_1_ receptors with losartan, we observed no attenuation of Ang-II-dependent modulation of SNr GABAergic neuronal APs with the specific AT_2_ receptor antagonist PD123319. Neither losartan nor PD123319 alone altered baseline AP firing characteristics in SNr GABAergic neurons. Additional angiotensin-related signaling modalities, such as other angiotensin molecules (e.g., IV and 1-7) and receptors (e.g., AT_4_ receptor and *Mas* receptors), could potentially participate in SNr GABAergic neuromodulation. Contributions by these mechanisms are likely minimal at best, given the relative abundance of Ang-II and AT_1_ receptors and our results with losartan blockade of AT_1_ receptors.

### Ang-II slows action potential kinetics and increases firing variability of SNr GABAergic neurons

Waveform analysis of Ang-II-responsive SNr GABAergic neuron APs revealed that Ang-II slows AP decay and rise, prolongs AP duration of evoked and spontaneous APs (except AP rise-time), and also promoted variability in the pattern of AP firing. These findings are consistent with reported changes in the electrical activity of several neurons exposed to Ang-II (Alioua et al., 2002; Zimmerman et al., 2005; Rainbow et al., 2009). Fast delayed rectifier Kv3.1 and Kv3.4 channels are primarily responsible for maintaining high sustained AP firing of SNr GABAergic neurons (Kaczmarek and Zhang, 2017; Ding et al., 2011; Zhou and Lee, 2011). Since AT_1_Rs reportedly inhibit delayed rectifier potassium channels in the hypothalamus and brain stem through a Gq-coupled protein kinase II-dependent pathway (Zhu et al., 1999), a similar mechanism may underlie the inhibitory effect of Ang-II on SNr GABAergic neurons. Future investigations are necessary to identify and mechanistically characterize ion channels potentially modulated by AT_1_ receptors in these cells.

### Ang-II activates postsynaptic GABA_A_ receptors in SNr GABAergic neurons

SNr GABAergic neurons receive robust inhibitory input from the striatum and external globus pallidus (Smith and Bolam, 1990; Lee et al., 2004; Deniau et al., 2007). Postsynaptic GABA_A_ receptors are the dominant inhibitory mechanism of these neurons (Precht and Yoshida, 1971; Boyes and Bolam, 2007)). Our data show that GABA_A_ receptor blockade with picrotoxin prevents Ang-II mediated suppression of evoked firing in SNr GABAergic neurons. We also observed an approximately 2.5-fold increase in picrotoxin-sensitive high-amplitude mIPSCs with Ang-II, but no noticeable change in average frequency and amplitude. This observation suggests that Ang-II activates postsynaptic GABA_A_ receptors in SN GABAergic neurons. Consistent with this hypothesis, Ang-II mediated facilitation of GABA_A_ receptor activity reportedly occurs in neurons of the rat median preoptic nucleus (Henry et al., 2009).

We also recorded SNr GABAergic mIPSCs following local Ang-II application; however, we observed no change in mIPSC properties. However, subsequent bath perfusion of Ang-II did increase the occurrence of high-amplitude mIPSCs. This observation suggests that GABA_A_ receptors potentiated by Ang-II are most likely expressed in distal dendritic terminals away from the soma and embedded deep in the slice. Further investigation is necessary to determine the location and mechanism underlying the coupling between AT_1_ receptors and GABA_A_ receptors in SNr GABAergic neurons.

### Ang-II does not change the action potential kinetics and basal IPSCs in SNc dopaminergic neurons

Evidence suggests that Ang-II via AT_1_ receptors impacts SNc dopaminergic neuronal function, homeostasis, and viability (Grammatopoulos et al., 2007; Garrido-Gil et al., 2013; Sonsalla et al., 2013; Labandeira-Garcia et al., 2014). At present, it is unclear if the effects of Ang-II on SNc dopaminergic neurons are direct, indirect, or a combination of both. To address this issue, we recorded evoked APs, PSCs, and IPSCs from SNc dopaminergic neurons. We could not reliably obtain interpretable AP firing frequency data due to the well-characterized adaptive response of dopaminergic neurons to depolarizing currents (data not shown). However, unlike SNr GABAergic neurons, we found that Ang-II had no apparent effect on the AP kinetics, PSCs, and IPSCs of SNc dopaminergic neurons, suggesting a heterogeneous effect in the two types of SN neurons.

The apparent lack of effect by Ang-II on SN dopaminergic neurons in our experiments could result from a low expression of GABA_A_ receptors, different subunit compositions, or a lack of canonical KCC2 K^+^-Cl^−^ co-transporters (Boyes and Bolam, 2007; Paladini and Tepper, 2016). SNr GABAergic projection neurons express KCC2 co-transporters, which actively pump [Cl^−^] out of the cell to maintain a hyperpolarized (around −71 mV) chloride reversal potential (Rivera et al., 1999). In contrast, the chloride reversal potential is around −63 mV in SNc dopaminergic neurons due to a lack of KCC2 expression (Gulácsi et al., 2003). In this scenario, GABA_A_ receptor potentiation could produce a greater hyperpolarization in SNr GABAergic neurons than SNc dopaminergic neurons.

### Ang-II decreases light-evoked EPSPs in SNr GABAergic neurons

During periods of synchronous activity, localized feedback inhibition within the SNr limits SNr GABAergic neuronal output (Brown et al., 2014). This inhibitory mechanism involves SNr GABAergic axon collaterals, which synapse not only with dopaminergic neurons in the SNc but also with SNc GABAergic neurons. The resultant collateral inhibition provides a robust gain control of the total GABAergic neuronal output (Tepper et al., 1995; Tepper and Lee, 2007; Brown et al., 2014).

Using transgenic mice with GABAergic neuron-specific expression of ChR2 (THY1-ChR2-YFP), we examined the negative modulatory effect of Ang-II on SNr GABAergic neurons during synchronous phasic activity. Consistent with our electrically-evoked experiments, we found that Ang-II decreased light-evoked EPSPs (intraburst suprathreshold spikes) in 4 out of 6 SNr GABAergic neurons. These data further confirm the inhibitory nature of Ang-II on SNr GABAergic neurons. Given the complex and non-uniform arborizations of the SNr GABAergic axon collaterals, we suggest that the effect of Ang-II on SNr GABAergic neurons is likely to be heterogeneous. An individual SNr GABAergic neuron’s response to Ang-II during phasic stimulation will depend on the extent of the local connections with neighboring GABAergic neurons. As such, the effect of Ang-II on SNr GABAergic output is complicated further by the influence of heterogenous intranigral feedback activity.

### Ang-II enhances feedforward inhibitory input to SNc dopaminergic neurons

SNr GABAergic neurons provide robust, monosynaptic inhibition of SNc dopaminergic neurons (Tepper et al., 1995; Tepper and Lee, 2007; Brazhnik et al., 2008; Pan et al., 2013). Ang-II-mediated suppression of SNr GABAergic output should, therefore, have a disinhibitory effect on SNc dopaminergic neurons. We observed, unexpectedly, that Ang-II increased the amplitude of SNc dopaminergic neuron IPSCs and IPSPs in response to photostimulation of SNr GABAergic neurons. This paradoxical observation is inconsistent with our other data showing Ang-II mediated suppression of SNr GABAergic neurons. This nonlinear effect of Ang-II on SNr GABAergic neurons and their output to postsynaptic SNc dopaminergic neurons highlights the complex nature of intranigral circuitry and neurotransmission.

SNr GABAergic projection neurons have extensive intra- and extranigral collateralizations. Non-uniformity of these branchings further contributes to variations in excitability, synaptic bouton content, and ion channel distributions within the network (Debanne et al., 2011; Martin et al., 2014; Seidl, 2014; Bucher, 2016). Such complexity is likely to be evident in the neuronal responses at the microcircuitry level within the SN. Indeed, the summed activity from a heterogeneous and interlinked network could result in a range of synchronous or asynchronous population activities. Therefore, the paradoxical facilitation of inhibitory input to SNc dopaminergic neurons by Ang-II upon phasic activation of SNr GABAergic neurons could be explained, at least in part, by the complex and asymmetrical nature of the intranigral microcircuitry.

## Conclusions

From these data, we conclude that Ang-II signaling occurs in the SNr GABAergic neurons via a postsynaptic AT_1_R-dependent mechanism. Our data shows enhancement of postsynaptic GABA_A_ receptors and a slowing of AP kinetics by Ang-II as contributing factors for the suppressive effect of Ang-II in SNr GABAergic neurons. However, the inhibitory effect of Ang-II on SNr GABAergic neurons did not translate to disinhibition of postsynaptic SNc dopaminergic neurons. Rather, we observed a paradoxical increase in postsynaptic inhibitory input. This observation implies a nonlinear effect of Ang-II on GABAergic neurotransmission on postsynaptic SNc dopaminergic neurons. We suggest this reflects the complex and non-uniform intranigral inhibitory microcircuit formed by the extensive arborization of SNr GABAergic axon collaterals. Further investigation into the microcircuit dynamics, underlying signaling cascades and effector proteins (e.g., ion channels), and the ensuing effects on SNc dopaminergic neurons by Ang-II, is warranted.

## ACKNOWLEDGEMENTS

G.C.A. is supported by NIH 5R01HD087347 and J.V. is supported by NIH 5R01EY029227. The authors thank Drs. Andrew R. Rau, Shane T. Hentges, Susan L. Tsunoda and Michael M. Tamkun for providing technical assistance, insightful ciritiques, and helpful feedback.

